# A new paradigm for leprosy diagnosis based on host gene expression Insights from leprosy lesions transcriptomics

**DOI:** 10.1101/2021.07.30.454441

**Authors:** Thyago Leal-Calvo, Charlotte Avanzi, Mayara Abud Mendes, Andrej Benjak, Philippe Busso, Roberta Olmo Pinheiro, Euzenir Nunes Sarno, Stewart T. Cole, Milton O. Moraes

## Abstract

Transcriptional profiling is a powerful tool to investigate and detect human diseases. In this study, we used bulk RNA-sequencing (RNA-Seq) to compare the transcriptomes in skin lesions of leprosy patients or controls affected by other dermal conditions such as granuloma annulare, a confounder for paucibacillary leprosy. We identified five genes capable of accurately distinguishing multibacillary and paucibacillary leprosy from other skin conditions. Indoleamine 2,3-dioxygenase 1 (*IDO1*) expression alone was highly discriminatory, followed by *TLR10*, *BLK*, *CD38*, and *SLAMF7*, whereas the *HS3ST2* and *CD40LG* mRNA separated multi- and paucibacillary leprosy. Finally, from the main differentially expressed genes (DEG) and enriched pathways, we conclude that paucibacillary disease is characterized by epithelioid transformation and granuloma formation, with an exacerbated cellular immune response, while multibacillary leprosy features epithelial-mesenchymal transition with phagocytic and lipid biogenesis patterns in the skin. These findings will help catalyze the development of better diagnostic tools and potential host-based therapeutic interventions. Finally, our data may help elucidate host-pathogen interplay driving disease clinical manifestations.

**Author Summary:** Despite effective treatment, leprosy is still a significant public health issue in more than 120 countries, with more than 200 000 new cases yearly. The disease is caused mainly by *Mycobacterium leprae*, a slow-growing bacillus still uncultivable in axenic media. This limitation has hampered basic research into host-pathogen interaction and the development of new diagnostic assays. Currently, leprosy is diagnosed clinically, with no standalone diagnostic assay accurate enough for all clinical forms. Here, we use RNA-seq transcriptome profiling in leprosy lesions and granuloma annulare to identify mRNA biomarkers with potential diagnostic applications. Also, we explored new pathways that can be useful in further understanding the host-pathogen interaction and how the bacteria bypass host immune defenses. We found that *IDO1*, a gene involved with tryptophan catabolism, is an excellent candidate for distinguishing leprosy lesions from other dermatoses. Additionally, we observed that a previous signature of keratinocyte development and cornification negatively correlates with epithelial-mesenchymal transition genes in the skin, suggesting new ways in which the pathogen may subvert its host to survive and spread throughout the body. Our study identifies new mRNA biomarkers that can improve leprosy diagnostics and describe new insights about host-pathogen interactions in human skin.

## Introduction

Leprosy is a chronic infectious disease caused mainly by the slow-growing intracellular pathogen *Mycobacterium leprae* that does not grow in axenic media. This bacterium resides preferentially in skin macrophages and Schwann cells in peripheral nerves, inducing dermatosis and/or neuritis. Patients can present several distinct clinical forms according to their immune response, histopathological characterization, and bacterial load. A localized tuberculoid form (TT) is characterized by low bacterial counts and a strong cellular immune response. Conversely, in the opposite lepromatous (LL) pole, a disseminated form, patients exhibit several lesions, a predominantly humoral response, and a high bacterial load in the tissues [1–3]. Borderline forms are classified according to their proximity to the poles. For operational and treatment purposes, leprosy is classified by the World Health Organization as paucibacillary (PB) or multibacillary (MB), based on the number of skin lesions, associated with nerve involvement or the bacilli detection in slit-skin smears [4].

Early and precise diagnosis is instrumental to leprosy control since delay in diagnosis leads to late multidrug therapy, higher disability risk, and continuing transmission, as highlighted by the 200,000 new cases consistently reported annually in the last 10 years [4,5]. However, bacteriological, immunological, genetics or molecular methods are not sufficient for specific diagnosis when used alone. Diagnosis most commonly relies on clinical evaluation, occasionally complemented with histopathological examination and bacterial counts, but these procedures are mostly performed in national reference centers [4,6].

Efforts have been deployed to improve leprosy diagnostics using cutting-edge technologies, such as molecular identification of *M. leprae*, serological tests for specific bacterial antigens, and quantification of host biomarkers in plasma or *in vitro* whole blood assays (WBA) [7–9]. Overall, all methods outperform standard clinical diagnosis and can compensate for the low accuracy in detecting PB patients [4,7,8,10–14]. Yet, until now such investigations involved comparing confirmed leprosy cases against healthy endemic controls, who are not representative of individuals with suspected leprosy. Here, other skin conditions represent a better comparator.

Identification of markers for early infection is hindered by our poor understanding of pathogenicity and the mechanism by which patients develop one or the other form of leprosy, and nerve injuries [15]. Gene expression signatures have been used as diagnostic tools for several illnesses, from infectious [10–12,14] and autoimmune diseases [16,17] to cancer [18–20]. Some signatures have already been approved for clinical use [12,21–23]. In leprosy, findings from past studies indicate the great potential of expression profiling for disease diagnosis [24–27]. Nonetheless, they were limited by the number of patients [28], or lacked proper epidemiological controls, such as differential diagnosis groups.

Here, we applied a combination of bulk RNA sequencing and quantitative validation by RT-qPCR on RNA extracted from skin biopsies of various leprosy forms and from non-leprosy patients to define a specific leprosy host signature applicable to diagnosis. Then, we explored gene expression patterns to improve our understanding of the immunopathogenic mechanisms towards leprosy polarization.

## Results

### Discrimination of leprosy *vs.* non-leprosy lesions based on mRNA expression

RNA sequencing was used for pinpointing host candidate genes capable of differentiating leprosy lesions from one of the commonest differential diagnoses of leprosy, granuloma annulare (GA), and from healthy skin. RNA from skin lesions of all leprosy clinical forms (n=33), plus GA (n=4) and healthy skin (n=5) were sequenced (S1 Table). Differentially expressed genes (DEG) in leprosy *vs.* non-leprosy (GA + healthy skin) samples resulted in 1160 DEG with a |log_2_FC| ≥ 1 and FDR ≤ 0.01, with 961 upregulated in leprosy forms compared to non-leprosy (Fig 1A-B and S2 Table). Exploratory hierarchical clustering of the DEG with |log_2_FC| ≥ 1 and FDR < 0.01 grouped all patients’ samples into roughly two clusters, except for two: one BL leprosy and one GA that clustered apart from samples with the same diagnosis (Fig 1C). Gene Ontology enrichment analysis of up-regulated genes in leprosy compared to non-leprosy showed enrichment for biological processes associated with leukocyte activation, T-cell activation, immune response, response to the bacterium, neutrophil degranulation, cell killing, cytokine secretion, purinergic receptor signaling pathway, and regulation of defense response to viruses by the host (Fig 1D and S3 Table).

**Fig 1.**
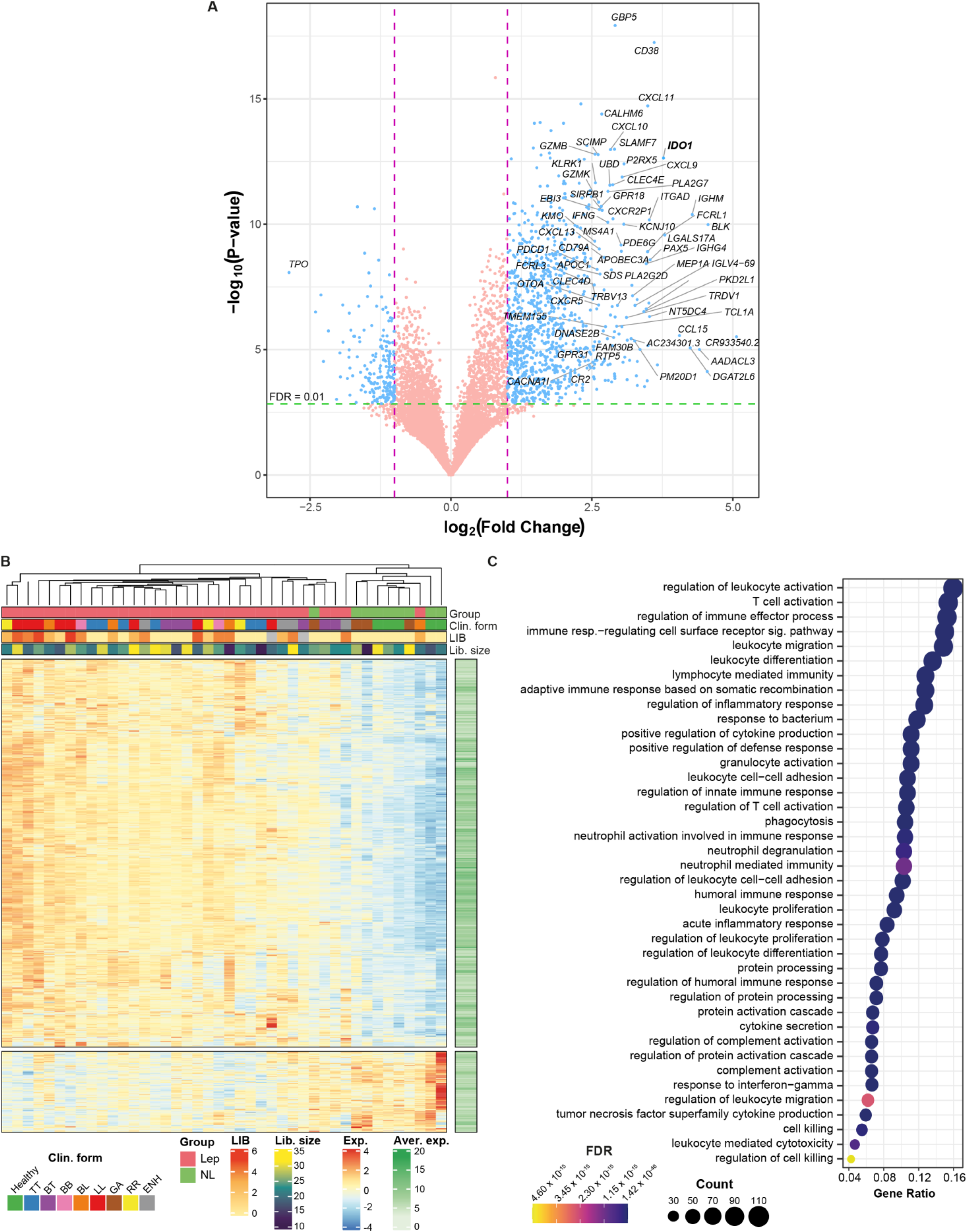
Differentially expressed genes from RNA-seq in leprosy *vs.* GA and leprosy *vs.* non-leprosy. (A) Volcano plot depicting DEG from leprosy *vs.* non-leprosy, where violet dashed line marks |log_2_FC| = 1. For clarity, gene symbols are shown only for the largest log_2_FC. (B) Heatmap with hierarchical clustering of samples based on expression of the DEG from leprosy *vs.* non-leprosy comparison. Color scale ranges from lower expression (blue) to higher expression (red). Library size is given in millions. LIB, logarithmic index of bacilli. (C) Biological processes from GO enriched for up-regulated DEG from leprosy *vs.* non-leprosy comparison. FDR, false discovery rate; NL, non-leprosy; GA, granuloma annulare; non-leprosy: GA + healthy individuals.

A total of 15 genes with the largest effect size (|log_2_FC| ≥ 1.5, FDR < 0.001), highest area under the curve (AUC), and plausible involvement with leprosy pathogenesis (S4 Table) were then validated using a two-step RT-qPCR with a new, larger, and more heterogeneous dataset including skin lesion samples from leprosy patients (n=25), and other common dermatoses (n=23) (S1 Table). Other dermatological diseases (ODD) included dermatitis (n=7), eczema (n=1), erythema (n=4), GA (n=6), lichen planus (n=2), psoriasis (n=2) and pityriasis alba (n=1) (S1 Table). A total of 12 samples per group was estimated to be sufficient to attain a power of 85% based on the Welch t-test (PB *vs.* ODD, MB *vs.* ODD) with alpha set at 0.03 to replicate the standardized effect size (log_2_FC/SD) estimated from RNA sequencing. Relative expression using the new sample set by RT-qPCR is shown in Fig 2A. Indeed, the validation data are in agreement with RNA sequencing, because 11 tested genes were replicated by RT-qPCR in terms of difference between mean expression (effect size in log_2_FC), except for *STAP1*, *GBP3*, *APOL3* and *CCR7* in PB *vs.* ODD comparison and *CCR7* in MB *vs.* ODD (Fig 2B-C, S5 Table). As for differentiating leprosy *per se vs.* ODD, genes *IDO1, BLK* (exon 11)*, CD38, CXCL11,* and *SLAMF7,* all had an area under the curve (AUC) of at least 96% with their lower bound 97% confidence intervals above 90% (Fig 2A, Fig 3C, S6 Table).

**Fig 2.**
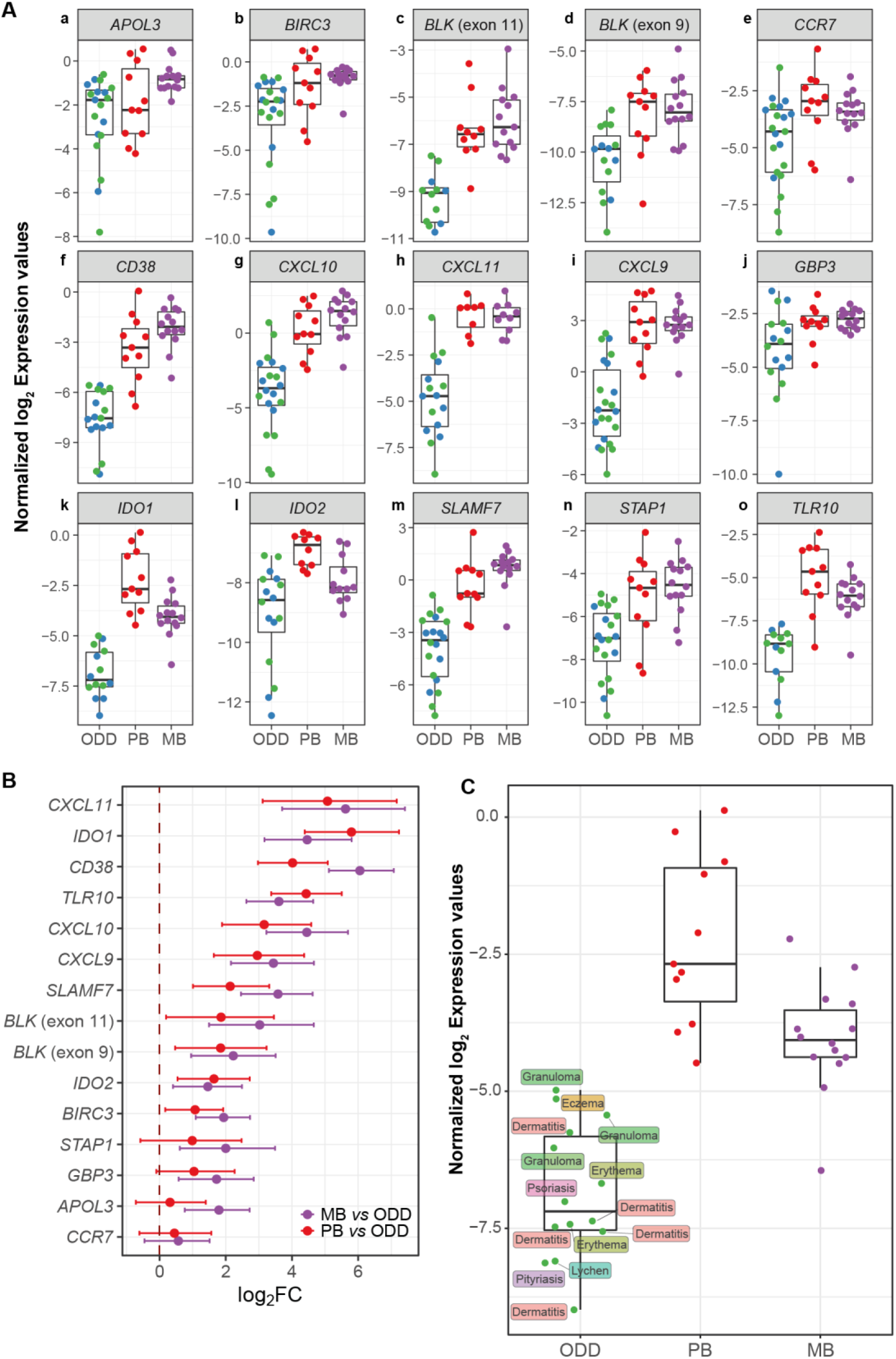
Technical and biological validation for selected DEG discovered from RNA sequencing. (A) Tukey boxplots with RT-qPCR normalized (2-3 reference genes) log_2_ expression values (A.U) according to clinical and histopathological diagnosis. ODD samples are colored according to *M. leprae* 16S rRNA qPCR status as positive (blue) or negative (green). (B) log_2_FC from MB-ODD and PB-ODD comparisons estimated from Bayesian linear mixed models and their 95% credible intervals. (C) Tukey boxplot highlighting *IDO1* RT-qPCR normalized log_2_ expression values by final diagnosis grouped into ODD category. Missing values are omitted.

**Fig 3.**
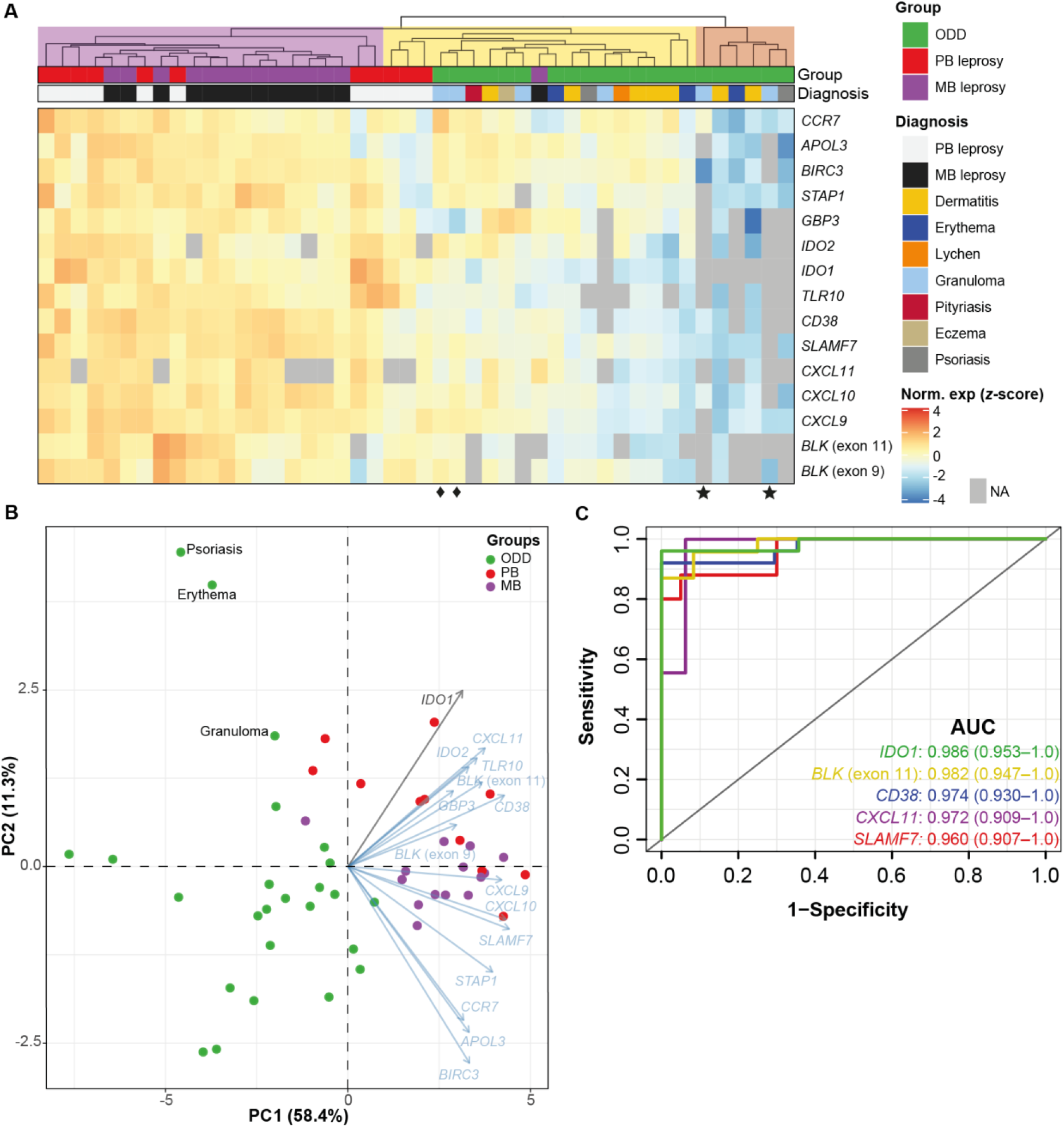
Hierarchical clustering of RT-qPCR replicated DEG and ROC analysis. (A) Hierarchical clustering with scaled and centered normalized log_2_ RT-qPCR expression values (arbitrary units) and annotated according to group and specific diagnosis. Dendrogram tree was cut arbitrarily and cluster analysis is for hypothesis generating purposes only. Two samples had more than 13 missing expression values and were removed from A. (B) Principal component analysis (PCA) with 15 genes measured by RT-qPCR and using log_2_ normalized scaled data. For PCA only, missing values were imputed by the gene arithmetic mean. NA, not amplified, i.e., Cp > 40. In this regard, there were two outliers (psoriasis and erythema), which are samples with high numbers of NA values and that were imputed using the gene arithmetic mean. (C) Receiver operating characteristic analysis for genes with largest AUC (97% confidence intervals) from RT-qPCR replication samples (complete data are shown in S6 Table). See also S1 Appendix and S1 Fig.

Next, hierarchical clustering with RT-qPCR data including missing values for some genes (no target gene amplification by RT-qPCR) was performed to examine all samples simultaneously. The analysis roughly revealed three major clusters (Fig 3A). At the highest tree subdivision, one small cluster (n=6) with the dendrogram grouped in light brown was composed of ODD samples with lower expression levels (Fig 3A). Due to several ODD having missing values, we confirmed that these samples had similar gene expression for the reference genes, thereby eliminating the possibility of insufficient cDNA input. Another cluster, grouped in the light purple dendrogram, included all MB and most PB samples (except four in light yellow dendrogram). GA samples displayed two patterns, the first with two samples showing undetectable *IDO1* expression (Fig 3A, bottom star symbols). The second set (n=4) is scattered among other ODD samples (Fig 3A). It can be seen that GA and PB samples show highly similar expression profiles for some genes (Fig 3A bottom diamond symbols), reinforcing the difficulty in clinically discriminating between these two conditions, and underlining the relevance of their inclusion in our comparisons [29–31].

Then, by applying principal component analysis (PCA) to the 15 gene signature obtained with the expanded sample panel tested by RT-qPCR, we uncovered two major patterns separating leprosy lesions from ODD (Fig 3B). As expected, MB samples appeared more homogeneous than PB and ODD samples, while the latter were more dispersed revealing heterogeneous expression patterns (Fig 3B).

Next, we quantified the individual classification potential of these genes in distinguishing leprosy from ODD using ROC analysis on RT-qPCR data. *IDO1* expression alone was found to be 98% accurate using an arbitrary threshold, followed by *BLK* (exon 11), *CD38, CXCL11,* and *SLAMF7* (Fig 3C and S6 Table). Finally, to confirm the causal link between mycobacteria and our gene-set, we evaluated the mRNA profiles induced by other live-mycobacteria using a public RNA-seq dataset [32]. We observed that most gene expression signatures, including *IDO1,* could be successfully replicated as induced by either *M. leprae* and/or other mycobacteria (Fig 1 in Appendix S1 and S7 Table). By contrast, some of the tested genes such as *BLK*, *CXCL9*, *MS4A1,* and *TLR10* were not differentially expressed in any of the *in vitro* assays with mycobacteria (Fig 1 in Appendix S1 and S7 Table).

### MB and PB gene expression profiling and mRNA-based classifier

To define a small subset of genes with high classificatory potential (i.e. with non-overlapping expression values) to distinguish MB from PB lesions, we performed a penalized logistic regression (LASSO) model with k-fold cross-validation trained on the public microarray dataset [24]. This dataset was chosen because of the higher number of PB/MB samples compared to our RNA-seq dataset. As a result, three genes with non-zero coefficients were selected by the cross-validated LASSO model: *HS3ST2, CD40LG,* and *CCR6*, but only the first two genes were most frequently (∼80%) selected across 10,000 bootstrapped samples within the training dataset (Fig 4A-B). The median misclassification error estimated by the resampling was about 4% (±5.4% median absolute deviation), ranging from 0% to 32% (Fig 4C). Instability assessment in the number of selected genes by LASSO (Fig 4D) showed that most iterations resulted in four non-zero genes (range, 1-20). The final model containing the three genes (*HS3ST2*, *CD40LG,* and *CCR6*) was evaluated on two test RNA-seq datasets: our dataset and the one from Montoya *et al.* including MB (n=9) and PB (n=6) groups [28]. Penalized logistic regression demonstrated an accuracy of 100% (lower 95% CIs: 86.8% and 78.2%, respectively) in classifying MB from PB samples in both test RNA-seq datasets; yet, the Brier score indicated a better performance in Montoya’s et al. dataset, probably due to a more homogenous sampling (Fig 4E-F). The *HS3ST2* gene was consistently more expressed in MB leprosy lesions compared to PB, whereas the opposite was observed for *CD40LG* (Fig 4E-H) and *CCR6* (S2 Fig). In both datasets, the combined expression levels of *HS3ST2* and *CD40LG* showed good discrimination between the two groups (Fig 4E-H). However, given the sample size and the bootstrapped estimates, it is not currently possible to exclude *CCR6* from the model without additional replication.

**Fig 4.**
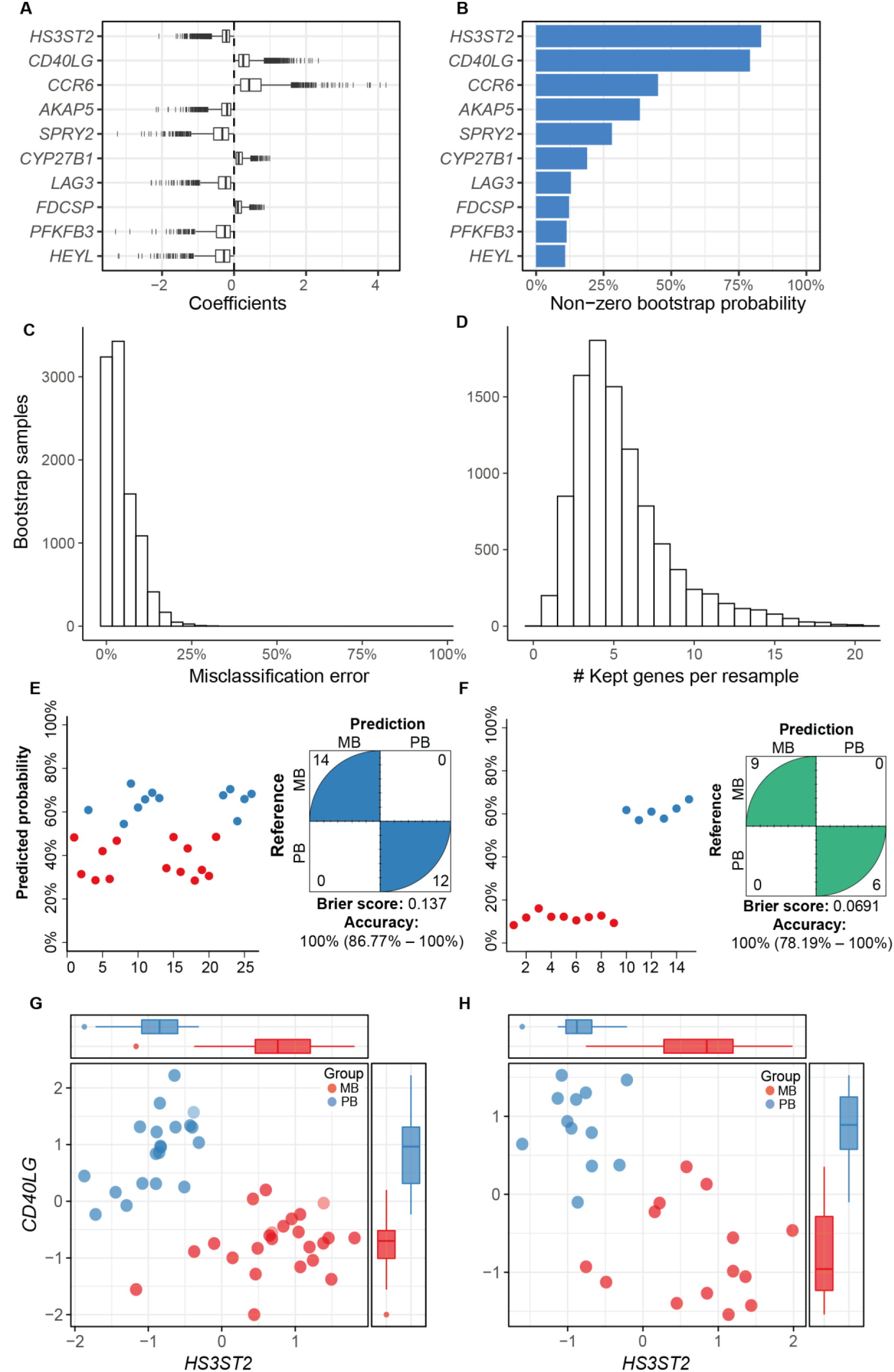
Gene candidates identified with the penalized logistic regression (LASSO) model as the most important to distinguish PB and MB leprosy lesions. (A) Coefficients (log odds) from the top 10 most selected genes (i.e., non-zero) across 10,000 bootstrap samples using the microarray from Belone *et al*. as training dataset. (B) Frequency of non-zero coefficients across all bootstrap samples. (C) Misclassification error distribution estimated from 4-fold cross-validation (k-) across 10,000 bootstrap samples, with median error of 3.70% (±5.4% median absolute deviation). (D) Number of genes kept across all resamples. Predicted probability from the final model performance on this study test RNA-seq (E) and Montoya *et al.* RNA-seq (F). Normalized log_2_ gene expression (z-score) of the two most frequently selected variables for distinguishing MB from PB samples in the (G) microarray training dataset and (H) this study test RNA-seq. PB, paucibacillary leprosy; MB, multibacillary leprosy. Tukey box plots with 1st, 2nd and 3rd quartiles ± 1.5 × inter quartile range (IQR) whiskers. See also S2 Fig.

Next, to assess the dichotomy beyond cellular *vs.* humoral response in leprosy lesions [33,34], a comparison of gene expression in MB leprosy (LL+BL+BB) *vs.* PB (TT+BT) skin lesions was performed. Differential expression analysis with |log_2_FC| ≥ 1 and FDR ≤ 0.01 resulted in 112 DEGs; 69 up-regulated and 43 down-regulated (Fig 5A and S8 Table). In addition, we compared DEG to the public microarray data available in Gene Expression Omnibus (GEO) from Belone *et al.* [24,35] using only the FDR cutoff. With an FDR < 0.01, 161 DEGs were common to both studies, all except one showed concordant modulation characterized by an overall high correlation coefficient and concordance index, irrespective of the technology used, the sample processing, and the data analysis methods (Fig 5B). Functional enrichment analysis of the RNA-seq up-regulated genes (i.e., more expressed in MB than PB) revealed processes involved with regulation of immune response, humoral immunity, phagocytosis, cholesterol metabolism, complement activation among others (Fig 5C and S9 Table). On the contrary, enrichment analysis of genes more expressed in PB revealed biological processes such as leukocyte differentiation, lymphocyte differentiation, lymphocyte-mediated immunity, B cell activation, STAT cascade activation/regulation, and JAK-STAT cascade activation (Fig 5D and S10 Table), which are consistent with exacerbated responses in granulomatous diseases. Localized clinical forms, i.e., BT and TT, show a gene expression pattern indicative of differentiation towards epithelioid transformation and granuloma assembly, which is also observed in cutaneous or pulmonary sarcoidosis [36,37].

**Fig 5.**
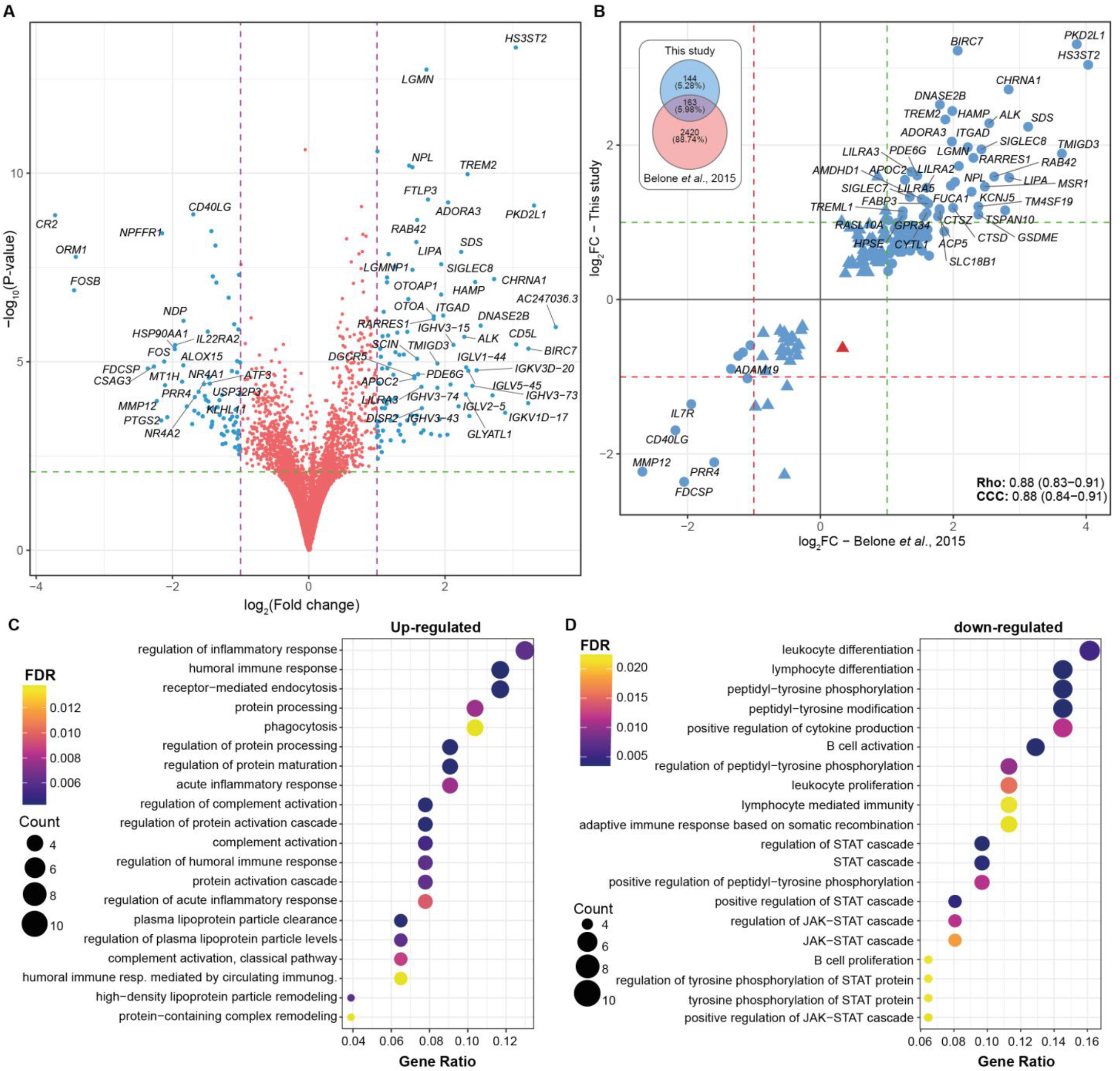
Differentially expressed genes from multibacillary (MB) *vs.* paucibacillary (PB) leprosy lesions. (A) Volcano plot showing DEG from the MB *vs.* PB comparison, where blue points are DE with |log_2_FC| ≥1 and FDR < 0.1. (B) Scatter plots with the 161 DEG common between this study and Belone *et al*. (24) microarray for the same comparison. Red and green dashed lines indicate log_2_FC of −1 and 1, respectively. Blue points are genes with the same modulation signal and red indicates discordancy. Rho, Spearman’s rank correlation coefficient. CCC, Lin’s concordance correlation coefficient. Venn diagram on the right displays the number of DEG in each study according to FDR < 0.01. (C) Biological processes from GO enriched from up-regulated and (D) down-regulated DEG. FDR, false discovery rate.

### Epithelial-mesenchymal transition (EMT) in the skin of multibacillary leprosy patients

To make the most of our dataset, we sought to test a previous hypothesis generated from our group’s microarray meta-analysis results, in which we have identified a consistent down-regulation of cornification, keratinocyte differentiation, and epidermal development-related genes in leprosy lesions, predominantly in MB [35]. We first hypothesized that such regulation could result from *M. leprae* inducing dedifferentiation of keratinocytes, similar to the phenomenon described previously in infected Schwann cells [38], and also seen in skin cancer by a process known as epithelial-mesenchymal transition (EMT) [39,40]. To test the hypothesis that such modulation was involved with EMT, we correlated the expression of the previously identified down-regulated genes in leprosy [35] with a collection of genes involved with previously Schwann cell dedifferentiation by *M. leprae* (Masaki *et al.* [38] signatures for EMT and non-EMT genes), positive markers of EMT (from literature), as well as annotated EMT and mesenchymal-related genes from Reactome (R.HSA.452723, R.HSA.5619507.3, R.HSA.2173791) and Gene Ontology (GO0001837) databases. Briefly, the normalized log_2_ expression matrices were filtered to retain only genes of interest. Then, the pairwise expression correlation for all genes was calculated using the Spearman’s rank correlation procedure. Finally, after adjusting the P-values for multiple testing, the genes with any pairwise correlation passing FDR ≤ 1 × 10^−4^ and *rho* ≤ −0.8 were visualized using a heat plot. As result, with this study’s RNA-seq, we found a consistent moderate negative correlation between keratinization, cornification, and epidermal development genes (Fig 6A, blue stars, *AQP3, DMKN, DSG1, DSP, EFNB2, JAG1, JAG2, KRT5, KRT10, KRT15, KRT19, OVOL2, PKP1, TACSTD2*) with those involved with canonical/alternative EMT and mesenchymal phenotypes (Fig 6A, green stars, *CTSZ, MMP9, PSAP, RHOA, TGFBR1, TGIF2, ZEB2, TGFB1*). Interestingly, the strongest correlations with epidermal/keratinocyte genes was with TGFβ-EMT-related genes (Fig. 6A blue block), as opposed to Masaki et al. non-EMT and other mesenchymal/pluripotency pathways. Next, we replicated these observations with Belone *et al.* microarray [24] and Montoya *et al.* RNA-seq datasets [28], respectively. In Fig 6BC the strongest and representative correlations from TGF□-EMT-related pathway and a keratinocyte/epidermal gene signature are shown in detail, while the remaining are available in Fig. S3-4.

**Fig 6.**
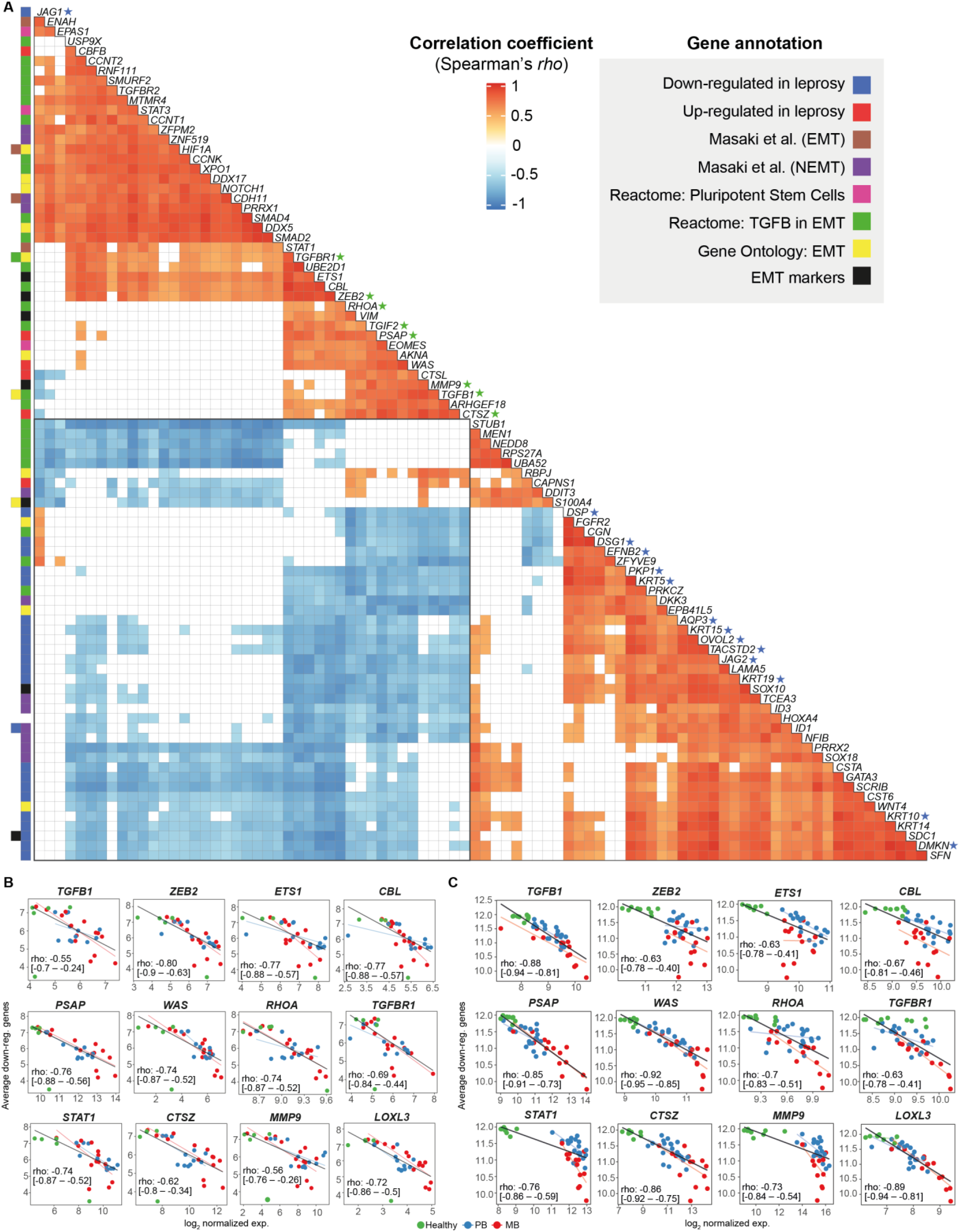
Strongest correlations between keratinocyte and EMT-related genes in leprosy lesions. (A) Heat plot with Spearman’s *rho* correlation coefficient of the strongest correlations after multiple testing adjustment with at least one gene-pair passing FDR ≤ 0.0001 and rho ≤ −0.8. Correlations with FDR > 0.1 are filled with white. Row colored squares identify gene annotations. Scatter plots of average log_2_ expression calculated with keratinocyte/epidermal development-related genes previously documented as down-regulated in leprosy skin against dedifferentiation-related genes using either (B) this study RNA-seq dataset or (C) Belone *et al.* microarray (GSE74481). Lines were drawn based on intercept and beta parameters estimated from robust linear regression for all samples (black line) or separately for PB (blue line), and MB (red line). Spearman’s *rho* coefficient along with 95% nominal confidence intervals are shown inside scatter plots calculated from all samples. See also S3 Fig and S4 Fig.

Overall, these results showed a decreased expression pattern of EMT-related genes in healthy skin samples, and a linear expression increase in PB and MB patients, especially with the microarray dataset, except for *MMP9* (Fig 6C). This was accompanied by the previously reduced expression of cytokeratins and epidermal development genes observed in leprosy. From these results, we hypothesize that in addition to TGF□-dependent immunosuppression in MB patients, activation of this pathway could be slowing or arresting keratinocyte cornification processes in leprosy lesions thereby both facilitating survival and/or spread of *M. leprae*. If not involved with dedifferentiation of keratinocytes or other epithelial cells, an alternative explanation would be loss of epithelial barrier in MB patients, possibly enlightening a new *M. leprae* transmission route. Further mechanistic experiments ought to determine the causality of our observations and test these findings in light of our hypothetical explanations of the phenomenon.

## Discussion

One of the priorities in leprosy research is the development of reliable and accurate laboratory diagnosis tools for all leprosy forms to provide efficient treatment and prevent disability [41]. This goal includes diagnosing patients with early forms of the disease, those with low or mild apparent symptoms, thus assisting with ambiguous differential diagnoses, and even classifying the disease for treatment (MB *vs.* PB) [4].

Host response to infection as measured by gene expression in skin biopsies offers diagnostic, prognostic and predictive potential. By applying host transcriptomics to skin lesions from leprosy patients and other common confounding dermatoses that challenge clinicians and pathologists [9,30], we identified a small set of genes that provide a promising expression signature capable of distinguishing PB leprosy cases from other confounding dermatological diseases. The top candidate, *IDO1,* is a gene involved in nutritional immunity and metabolism [42–45]. Alone, the expression of this gene was able to differentiate leprosy from non-leprosy lesions with high accuracy in our dataset and in others. According to the latest data from single-cell analysis [46], *IDO1* has been shown to be differentially expressed in Langerhans cells from leprosy lesions compared to healthy skin, corroborating our findings. However, *IDO1* expression is also increased in other mycobacterial diseases such as tuberculosis [47,48], which might decrease its specificity. The accuracy of classification could be improved by combining measurement of *IDO1* expression with that of four other biomarker genes *BLK, CXCL11, CD38, TLR10 and SLAMF7,* which also showed high classification accuracy in the replication dataset. In parallel, the penalized logistic regression model, evaluated on two independent datasets, demonstrated that *HS3ST2* and *CD40LG* hold potential to differentiate between MB and PB lesions. In parallel, the penalized logistic regression model, evaluated on two independent datasets, demonstrated that *HS3ST2* and *CD40LG* hold potential to differentiate between MB and PB lesions. We recognize that there is no clinical utility in classifying MB from PB lesions with laboratory assays because this can be done during anamnesis alone. Hence, we aimed at identifying molecular features differing not only in the measure of effect (log_2_FC) but also having little overlap between the lesion types, as this may point to previously unexplored genes and pathways relevant to future investigation. Considering the functional evidence for *HS3ST2* [49], it is possible that this gene may be involved with granuloma disassembly, tissue permeability, and cellular migration in leprosy, which would explain its overexpression in MB lesions. On the contrary, *CD40LG* (also known as CD154) is more expressed in PB patients when compared to MB with a predominant role in the activation of the microbicidal *Th1* response associated with PB lesions [50]. After mechanistic validation of our findings, quantifying expression levels of *HS3ST2* and *CD40LG* from leprosy lesions could be useful to assess immune responsiveness against *M. leprae*, help patient stratification and/or provide a basis for host-based adjuvant treatment for leprosy lesions.

One of the challenges in translating gene expression signatures into medical diagnosis is the cost of measuring a large number of genes and transforming these values into a unique continuous or binary classifier. So far, we were able to reproduce the findings using both bulk RNA-sequencing and relative RT-qPCR, with the latter being more accessible to clinicians at least in reference centers or central hospitals. Although there are successful approved RT-qPCR relative gene expression-based diagnostic tests for diagnosing sepsis [12], clinical support for prostate [22], and breast cancer [18], there is a need for alternatives to reduce the cost and complexity of such assays. Quantification of mRNA based on isothermal amplification either with NASBA [51,52], RT-LAMP [53,54] or CRISPR-Cas12 [55] is conceivable for less specialized settings without high-end equipment. Besides, combining a multi-target expression-based diagnostic test with qPCR detection of *M. leprae* DNA could increase the specificity and sensitivity of leprosy diagnosis [56]. Alternatively, an ELISA assay measuring the levels of IDO1 protein from skin interstitial fluid, for example, could be proven useful [57]. Further studies ought to be done selecting tangible diagnostic thresholds and devising a proper classification system to allow the biomarker to function unsupervised.

In parallel with poor diagnosis, lack of fundamental understanding of leprosy pathogenesis has misled scientists for centuries [5,6]. Herein, we also compared the two leprosy poles, MB and PB, and identified several pathways already known to be associated with leprosy, such as the humoral immune response, phagocytosis, and complement activation. Genes involved with cholesterol and fatty acids were more expressed in MB lesions, as already reported [58–60]. Interestingly, B-cell-related genes were more expressed in PB than MB. In fact, it seems that both poles modulate this pathway by a distinct set of genes. Involvement of B lymphocytes in PB leprosy pathogenesis has been described by a few groups, which may indicate differential involvement of such cells depending on the disease pole [61,62].

*M. leprae* subverts host cell metabolism [63] by inducing lipid biosynthesis, while avoiding type II (IFN-gamma) responses through a type I IFNs mechanism, following the phagolysosomal breach that releases DNA into the cytosol [64]. However, exactly how the bacilli spread throughout the body and bypass the microbicidal immune response remains unknown. Here, we provide robust evidence indicating that *M. leprae* may induce EMT in the skin within keratinocytes and macrophages, as described in Schwann cells [38]. Indeed, *M. leprae* induced dedifferentiation of infected Schwann cells into an immature stage resembling progenitor/stem-like phenotype [38]. These reprogramming events induced by long-term infection with *M. leprae* resulted in mesenchymal cells capable of migratory and immune-permissive behavior, which in turn facilitated *M. leprae* spread to skeletal and smooth muscles and furthered macrophage recruitment [38,65]. In our previous work, we identified a down-regulated signature of keratinocyte differentiation and cornification gene markers in MB skin lesions [35]. Here, we showed that such genes are inversely correlated with genes involved with EMT, especially the members of the TGFβ-EMT pathway, such as *TGFB1*, *TGFBR1*, *TGIF2, PSAP, ZEB2* [66,67]. Some of these genes are directly or indirectly associated with EMT, such as a *PSAP* [68], *WAS* [69], *RHOA* [70–73], *CTSZ* [74], *MMP9* [75], *LOXL3* [76], *HIF1A* [77,78] among others.

Our hypothesis that *M. leprae* is inducing dedifferentiation or slowing the cornification process in keratinocytes is plausible, given the evidence in Schwann cells and a few reports of infection in this cell type (Fig 7) [79,80]. Nevertheless, other phenomena could explain EMT’s role in leprosy pathogenesis, such as wound healing or loss of the epithelial barrier. Although, given its obligatory intracellular lifestyle, *M. leprae* induces dedifferentiation in other cell types, either directly as in Schwann cells or indirectly via chemokine and cytokine production in lesions. Besides inducing keratinocyte dedifferentiation to mesenchymal cells, *M. leprae* might benefit from a decreased or alternative immune activation of these cells [81,82]. Further functional confirmatory experiments should elucidate the causality of this correlation and provide definitive evidence of the relationship between the bacilli and other cell types, such as keratinocytes, fibroblasts, and epithelial cells.

**Fig 7.**
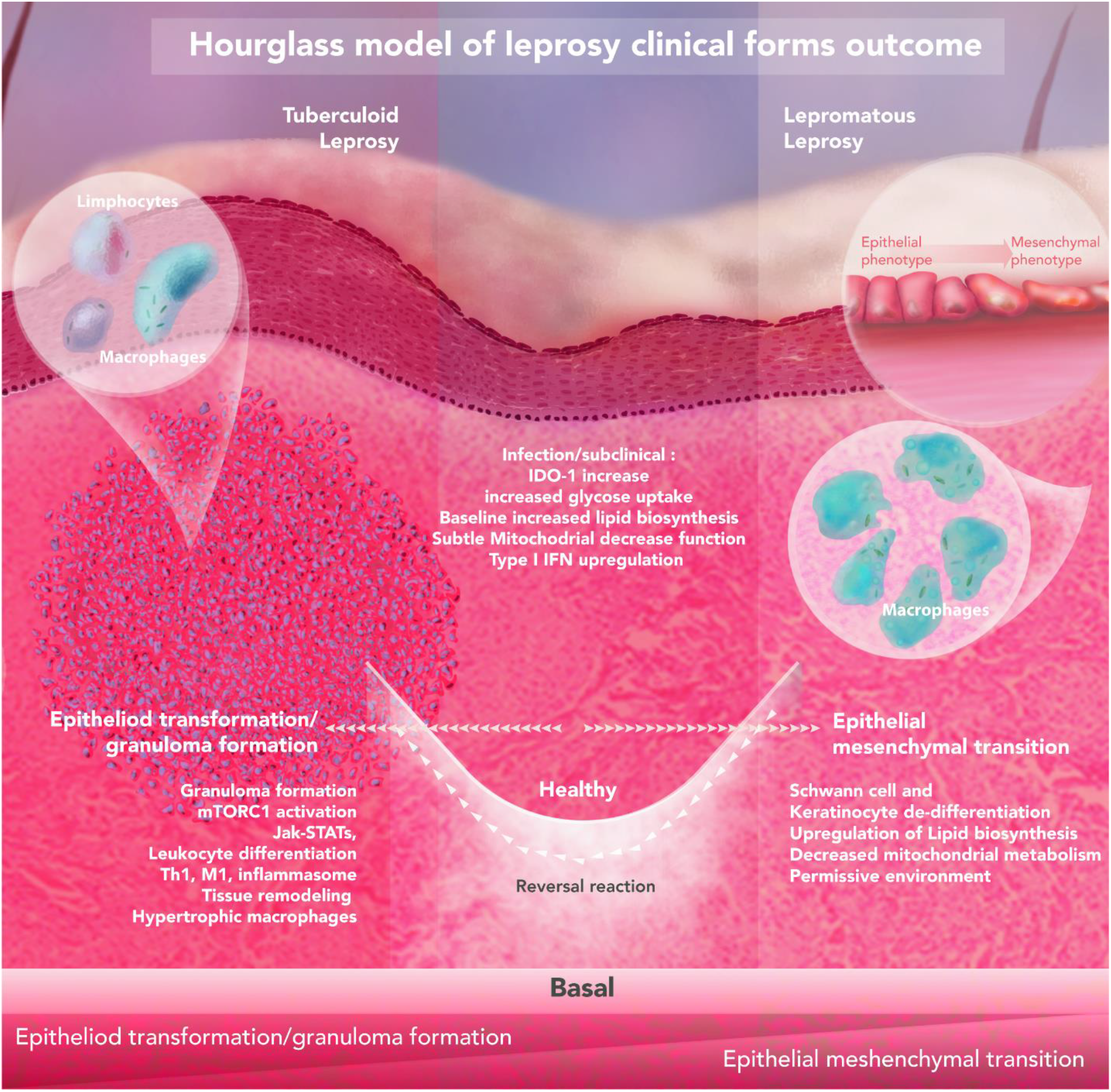
Hypothetical hourglass model contextualizing the observed findings for leprosy clinical outcomes. The host-pathogen interaction in the skin leads to opposing leprosy clinical forms. Upon infection, *M. leprae* induces baseline metabolic alterations such as an increase in glucose uptake, modulation of lipid biosynthesis, reduction of mitochondrial metabolism, and upregulation of IDO-1 and type I IFN. Eventually, progression towards an unspecified inflammatory state can be observed where three ways could be anticipated: I) self-healing; II) progression towards the tuberculoid pole; or III) progression to lepromatous pole. These outcomes are driven by specific environmental and host genetic factors. It is expected that lower (or shorter) *M. leprae* exposure, food shortage, BCG vaccination, and polymorphisms in genes controlling autophagy/granuloma formation (*NOD2*, *LRRK2*, *PRKN*) all contribute to developing leprosy per se. Excessive inflammation is one phenotype observed, that is also seen in other granulomatous diseases (e.g., cutaneous sarcoidosis, granuloma annulare), especially in paucibacillary lesions. On the other pole, epithelial-mesenchymal transition and local immunosuppression are present due to a probably higher (and/or longer) *M. leprae* exposure, combined with host single-nucleotide polymorphisms (SNPs) at key genes, like lipid biogenesis (*APOE*) and central metabolism (*HIF1A, LACC1/FAMIN*), culminating in disease progression.

Our preliminary data also showed that the enriched pathways among PB skin lesions were consistent with profiles observed in other granulomatous diseases, such as noninfectious sarcoidosis and granuloma annulare, or chronic infectious diseases like tuberculosis [37,83–85]. Our findings revealed that PB (TT/BT) lesions have, among others, JAK-STAT cascade activation, which has been implicated in sarcoidosis and GA. Remarkably, the JAK-STAT specific biological inhibitor, tofacitinib, has a potent effect promoting rebalance of exacerbated immunity among sarcoidosis and granuloma annulare patients reestablishing homeostasis [83]. Another compound, everolimus, has been shown in experimental models to achieve the same response [37] suggesting that these drugs could be useful to treat PB, but not MB, leprosy.

To conclude, our combined findings provide highly discriminatory mRNA signatures from skin lesions that could distinguish leprosy from other dermatological diseases and allow disease classification by monitoring only a handful of genes. In addition, we report new genes and pathways that are likely informative regarding how *M. leprae* interacts with and subverts host cells to promote its spread within the body and subsequent transmission.

## Materials and Methods

### Patient cohort

All patients were enrolled after informed written consent was obtained with approval from the Ethics Committee of the Oswaldo Cruz Foundation, number 151/01. Leprosy clinical forms were classified according to the criteria of Ridley and Jopling [2]. Leprosy patients were treated according to the operational criteria established by the World Health Organization [4]. Leprosy and patients with other dermatological diseases were eligible if their diagnosis was confirmed by clinical and histopathological findings. Additionally, detection of *M. leprae* DNA by qPCR routinely performed in our laboratory could be employed to support diagnosis [56,86]. HIV and hepatitis B positive patients were not included in this study, in addition, we excluded individuals with a current or previous history of tuberculosis. No other comorbidities were used to exclude patients and further individual information is available in S1 Table. Skin biopsy specimens containing both epidermis and dermis were obtained with 3 mm (diameter) sterile punches following local anesthesia from the lesion site. Skin biopsies were immediately stored in one milliliter of RNALater (Ambion, Thermo Fisher Scientific Inc., MA, USA) according to the manufacturer’s instructions and stored in liquid nitrogen until RNA isolation. Healthy skin biopsies were from lesion-free sites of patients diagnosed with indeterminate or pure neural leprosy.

### Study Design

The main objective of this research was to identify host gene expression patterns capable of distinguishing leprosy (including the PB forms) from other differential diagnosis of skin lesions. Our working hypothesis was that leprosy lesions, despite their morphological and histopathological similarity to other skin diseases, may induce distinct patterns of gene expression in at a small subset. We predefined the comparison of leprosy (PB+MB) from non-leprosy including GA in addition to healthy patients for RNA sequencing experiment. In addition, we predetermined comparisons between leprosy poles: MB *vs.* PB. Our samples are representative of a population of individuals attending the Sousa Araujo Outpatient Clinic based in Rio de Janeiro, Brazil, which also receives patients from surrounding municipalities.

### RNA isolation

Snap frozen skin biopsies were thawed in wet ice and processed using TRIzol Reagent (Ambion, Thermo Fisher Scientific Inc., MA, USA) according to the manufacturer’s instructions with the help of Polytron Homogenizer PT3100 (Kinematica AG, Switzerland). RNA was treated with DNAse using the DNAfree kit (Thermo Fisher Scientific Inc., MA, USA) according to the standard manufacturer’s protocol, prior to use for library preparation and RT-qPCR. RNA integrity was assessed in 1% agarose gel electrophoresis or TapeStation RNA ScreenTape (Agilent Technology, CA, USA). During RNA isolation, samples were randomly assigned to extraction batches and freeze-thaw cycles to minimize batch effects and the introduction of technical artifacts. All procedures applied to samples were carried out using reagents from the same lot. The first author conducted the experiments aware of each sample group during the entire process, therefore, no blinding scheme was used, although we do not rely on perceptual/abstract measurements or analyses nor did we purposefully exclude samples.

### Library preparation and Illumina RNA sequencing

RNA-seq libraries were prepared with 1 µg of total RNA for each sample using the Illumina TruSeq mRNA kit (Illumina, USA) as recommended by the manufacturer using the Illumina CD RNA indexes (Illumina, USA). Libraries were quantified and qualified using a qPCR quantification protocol guide (KAPA Library Quantification Kits for Illumina Sequencing platforms) and TapeStation D1000 ScreenTape (Agilent Technologies, USA), respectively. The resulting libraries (fragment size 200-350bp) were multiplexed (17, 17, and 19 libraries, respectively) and sequenced using the NextSeq 500 platform (Illumina, USA), generating approximately 520 million single-end reads of 75 nucleotides in length.

### RNA-sequencing analysis

RAW bcl files were converted into .fastq using Illumina’s bcl2fastq script. Then, read quality was assessed using FastQC version 0.11.8 [87]. Next, transcript counts were estimated using Salmon (v.1.13.0) quasi-mapping (human transcriptome GRCh38_cdna sourced from Ensembl/RefGenie plus pre-computed salmon index, http://refgenomes.databio.org/#hg38_cdna) with default settings and --seqBias flag set [88]. Transcript counts were summarized into ENSEMBL gene counts using the R v.3.6.1 package tximport v.1.12.0 [89,90] and biomaRt v.2.40.5 [91]. The expression of sex-chromosome-specific genes, such as *UTY* and *XIST,* was used to rule out sample mislabeling. Differential expression was estimated using DESEq2 v.1.24.0, after filtering out weakly expressed genes with less than 10 counts per million and less than 15 total counts in 70% of samples [92–94]. In addition to the patient’s biological sex, extraction batch and sequencing run, three surrogate variables estimated with RUVseq v.1.18.0 were included in DESeq2’s generalized linear model [95,96]. Nominal P-values were inspected with histograms and adjusted for multiple testing according to the method [97] proposed for controlling the false discovery rate (FDR). All log_2_ fold-changes were shrunken prior to DE filtering with the apeglm [94] or normal algorithms. For visualization, counts per million (CPM) were computed with edgeR’s cpm function v.3.26.1 and variance stabilized with the parametric method [92]. Then, surrogate variables and covariates were regressed out from the expression matrix using limma’s removeBatchEffect [98–100] before being visualized with ggplot2 v.3.3.0 [101]. Hierarchical clustering, heatmaps, and ROC analysis were all performed with the previously processed expression matrix. Heatmap with hierarchical clustering was drawn with ComplexHeatmap v.2.0.0 [102] or pheatmap v.1.0.12 [103] using gene-wise scaled and centered matrix with Euclidean distance and average agglomeration method. Overrepresentation analysis (ORA) was used to test for Gene Ontology Biological Process (GO BP) enrichment with clusterProfiler v.3.12.0 [104] and org.Hs.eg.db v.3.8.2 annotations [105]. Up and down-regulated lists were used as inputs and the background list was composed of all genes subjected to differential expression. P-values were adjusted for multiple testing using the Benjamini-Hochberg method [97]. Raw and normalized RNA sequencing data are available in EMBL-EBI’s ENA and ArrayExpress under accessions ERP128243 and E-MTAB-10318, respectively.

### RT-qPCR

A total of 2.5 µg of RNA was reversed transcribed into cDNA using 4 µL of Vilo Master Mix (Thermo Fisher Scientific Inc., USA) according to the manufacturer’s instructions. Then, cDNA was diluted to a final concentration of 5 ng/µL using TE buffer (10 mM Tris-HCL and 0.1 mM EDTA in RNAse-free water). RT-qPCR was performed using Fast Sybr Master Mix (Thermo Fisher Scientific Inc., USA) in a final reaction volume of 10 µL. For each reaction, performed in duplicate, 5 µL of Fast Sybr Green were combined with 200 nM of each primer, 10 ng of cDNA, and q.s.p of injection-grade water. Thermal cycling and data acquisition were performed on Viia7 with 384 well block (Applied Biosystems, Thermo Fisher Scientific Inc., USA) following the master mix manufacturer cycling preset with a final melting curve analysis (65 °C to 95 °C, captured at every 0.5 °C). All primers were designed with NCBI Primer-Blast [106–109] to either flank intron(s) or span exon-exon junction(s) to avoid gDNA amplification (S11 Table). Further, primers were quality checked for specificity, dimers and hairpin with MFEPrimer v.3.0 [110,111] and IDT’s oligoAnalyzer (https://www.idtdna.com/calc/analyzer). Data were exported from QuantStudio software v.1.3 in RDML format, which was imported to LinRegPCR v.2020.0 for RT-qPCR efficiency determination and calculation of the N_0_ value [112,113]. Finally, N_0_ values were imported to R and normalized using as the denominator the normalization factor (NF) calculated from the geometric mean of at least three reference genes (*RPS16*, *RPL35* and *QRICH1*), which were previously tested for stability [114]. These N_0_ normalized values were used for visualization in Fig 2A. For mean difference estimation between groups, RT-qPCR data were analyzed in a Bayesian framework (Markov Chain Monte Carlo sampling, MCMC) using generalized linear mixed effect models under lognormal-Poisson error with MCMC.qpcr v.1.2.4 [115,116]. Per-gene efficiency estimates from LinRegPCR were used in conjunction with Cp (crossing point) calculated in QuantStudio software v.1.3 to generate the counts table. Then, the generalized linear mixed-effect model was fitted using three reference genes (allowing up to 20% between-group variation) with 550,000 iterations, thin = 100, and burn-in of 50,000. The model specification included the sample (factor with 51 levels) as a random effect and the diagnosis group (factor with 3 levels) as a fixed effect. MCMC diagnostics were done by inspecting chain mixing plots and linear mixed model diagnostic plots. Ninety-five percent credible intervals were drawn around the posterior means and MCMC equivalent P-values were also computed.

### Reanalysis of public gene expression datasets

Belone and collaborators GSE74481 [24] and de Toledo-Pinto and cols. GSE35423 [64] microarray datasets were reanalyzed as described elsewhere [35]. Blischak and cols. [32] RNA-seq dataset (GSE67427) was reanalyzed from counts per sample file from the author’s Bitbucket repository (https://bitbucket.org/jdblischak/tb-data/src/master/). Briefly, a normalized log_2_ expression matrix was regressed out for RNA integrity number and extraction batch variables. Then, differences in gene expression (48h post-infection) for specific genes and treatments were tested using a gene-wise linear mixed model with a random intercept per sample (replicate) followed by Dunnet comparison against a “mock” group using emmeans v.1.5.3. Montoya and collaborators’ dataset was retrieved from GEO (GSE125943) already normalized (DESeq2 median ratio method) and transformed with base 2 logarithm with no further processing [28].

### Correlation analyses

For RNA-seq datasets, normalized log_2_ counts-per-million values were used and log_2_ normalized intensities for microarray. Spearman’s rank correlation method was chosen because it is robust against outliers, does not rely on normality assumption, and also identifies monotonic but non-linear relationships. Initially, a list of keratinocyte/cornification/epidermal development genes that were DE in the meta-analysis was assembled [35]. Then, lists of target genes were compiled from results of Masaki *et al.* [38]: EMT and non-EMT; from Reactome: R-HSA-452723 (Transcriptional regulation of pluripotent stem cells), R-HAS-5619507.3 (Activation of HOX genes during differentiation), R-HAS-2173791 (TGFβ receptor signaling in EMT); Gene Ontology GO:0001837 (EMT), and literature for EMT canonical markers. Next pairwise Spearman correlation was calculated using the Hmisc’s rcorr function v.4.2-0 for every pair of genes from keratinocyte/epidermal development and EMT gene lists. P-values were adjusted for multiple testing using the BH method for FDR control for all tests [97]. Additionally, 95% nominal confidence intervals were calculated using the Fieller method implemented by correlation R package v.0.5.0 [117,118]. To visualize the results, only genes with at least one pairwise correlation with Spearman’s rho coefficient ≤ −0.8 and FDR ≤ 0.0001 were selected. Additionally, the average log_2_ expression from genes involved with keratinocyte/epidermal development was calculated and used in scatter plots against the expression of the EMT genes. Scatter plots were drawn with ggplot2 v.3.3.3 showing lines from coefficients estimated using default robust regression (MASS::rlm v.7.3-51.4) either for all samples or stratified by group. No outliers were omitted.

### Regularized (LASSO) logistic regression classification

Normalized log_2_ expression matrices regressed out for covariates and batches were used as input predictors. The model was trained using the microarray dataset from Belone et al. [24] with penalized regression (L1-norm, LASSO) and 4-fold cross-validation (k-fold CV) with the negative binomial log-likelihood link function, glmnet v.4.1 [119–121]. Predictors were standardized to have mean zero and unit variance inside the cv.glmnet function. We opted for L1-norm because it results in a smaller number of genes (#features ≤ n) with non-zero coefficients, as compared to elastic-net or ridge regression counterparts. In addition, this model is suitable for high-dimensional data as it combines feature selection during model tuning and training, mitigating the effects of predictors’ collinearity and reducing overfitting. To assess the coefficients’ error, misclassification error rate, feature stability and model size we used non-parametric bootstrap (boot v.1.3.25) with 10,000 samples, with 4-fold cross-validation inside each loop [122,123]. The final LASSO model selected by 4-fold cross-validation contained three non-zero genes. Finally, independent RNA-seq test datasets were used to compute the accuracy of the final model. Alternatively, the whole process was repeated with leave-one-out cross-validation instead of k-fold. The results were practically indistinguishable, especially regarding the feature stability (data not shown).

### Sample sizes

The sample size for RNA sequencing was selected based on previous leprosy work with microarrays, aiming at detecting genes with at least a differential fold-change of two. For RT-qPCR validation, sample size calculation was performed using the per-gene standardized effect size estimated from the RNA-seq data, aiming at a power of 85% and alpha = 0.03. No samples were discarded after successful data collection (i.e. outliers). In the end, the sample sizes per group for RT-qPCR were: MB = 14, PB=11, ODD = 23. All RT-qPCR reactions were conducted in duplicate for each biological unit (here, a fragment of a skin biopsy derived from an individual).

### RT-qPCR and ROC statistical analyses

Normalized RT-qPCR gene expression data were log_2_ transformed before use in data visualization. Additionally, we checked if the Bayesian results remained consistent using a more common procedure (data not shown). For this, the mean normalized expression (from N_0_) was compared pairwise for the prior stipulated groups using Welch’s t-test implemented in R language, using the predetermined alpha of 0.03. Normality assumption was verified with normal quantile-quantile plots (qqplots, car v. 3.0-2). In cases where quantile-quantile plots showed huge deviation from theoretical normal distribution, the Wilcoxon Rank Sum was used to verify results.

Receiver Operating Curve (ROC) analysis was used to determine the accuracy (measured by the area under the curve, AUC) and its respective best classification threshold, aiming at maximizing AUC with equal importance for sensitivity and specificity. Confidence intervals (95%) for AUC were calculated using the Delong non-parametric method as implemented in pROC v.1.15.3 [124–126].

### Data and code reporting

Raw .fastq data are available in EMBL-EBI European Nucleotide Archive (ENA) database (ERP128243). Raw Salmon counts and normalized batch cleaned expression matrices are available in EMBL-EBI ArrayExpress, under E-MTAB-10318, along with experimental and phenotypic metadata. R source code and accompanying intermediate data used in all analyses in this manuscript are also readily available through Zenodo, doi.org/10.5281/zenodo.4682010.

## Acknowledgements

The authors wish to acknowledge Suelen Justo Moreira (MSc) and Rhana Prata (PhD) for assistance with skin biopsy RNA isolation. Helen Ferreira (MSc), Cristiane Domingues and José Augusto for their technical and logistic support. The Gene Expression Core Facility (GECF) at EPFL, Lausanne, Switzerland, especially Drs. Elisa Cora and Bastien Mangeat for sequencing assistance. All patients and staff (physicians, nurses and technicians) from Sousa Araujo Outpatient clinic at FIOCRUZ, Rio de Janeiro, Brazil.

## Supporting Information

**S1 Appendix. Linking expression profiles to mycobacteria species.**

**S1 Fig. Gene expression in MB and PB groups from test and training datasets.** Normalized log_2_ expression values per group from (A) this study RNA-seq dataset or (B) Belone *et al.* (GSE74481) [24]. The genes shown were selected in 25%–50% of the LASSO models (Fig 4B) according to the bootstrap. MB, multibacillary leprosy; PB, paucibacillary leprosy; TT, tuberculoid leprosy; BT, borderline-tuberculoid; BB, borderline-borderline; BL, borderline-lepromatous; LL, lepromatous. Each point represents an independent skin biopsy from a patient. Y-axis values are not comparable between panels A and B.

**S2 Fig. Strongest correlations between the average expression of genes associated with keratinocyte/cornification against dedifferentiation-related genes using Montoya *et al.* RNA-seq dataset** [28]. Scatter plots of scores (average normalized log_2_ expression) calculated from genes with previously documented down-regulation in leprosy skin lesions against dedifferentiation-related genes with Montoya *et al.* RNA-seq dataset (GSE125943) [28]. Lines were drawn based on intercept and beta estimates from robust linear regression for all samples (black) or separately for TL (tuberculoid leprosy, blue), and LL (lepromatous leprosy, red). X-axis shows log_2_ normalized expression values. Spearman’s rho are shown along with nominal 95% confidence intervals inside the plots. Most genes shown have FDR < 0.1 and rho ≤ - 0.6. Related to figure 6.

**S3 Fig. Strongest correlations between modulated genes from keratinocyte/cornification and dedifferentiation-related genes using Belone et al. microarray dataset (GSE74481)** [24]. Heat plot with Spearman’s rho correlation coefficient of the strongest correlations from all ontologies screened after multiple testing adjustment (BH-FDR). Most genes shown have FDR ≤ 0.0001 and rho ≤ −0.7. Bottom colored rectangles indicate which category the gene was present (some genes co-occur). Related to figure 6.

**S1 Table. Demographic and clinical metadata from human participants.**

**S2 Table. Genes differentially expressed from leprosy *vs.* non-leprosy with |log_2_FC| ≥ 1 and FDR ≤ 0.01.**

**S3 Table. Over-representation analysis (ORA) for leprosy *vs.* non-leprosy (up-regulated) differentially expressed genes.**

**S4 Table. ROC analysis from RNA-seq dataset using leprosy *vs.* non-leprosy samples.**

**S5 Table. Posterior log_2_FC estimates, 95% credible intervals and MCMC P-values from PB-OD and MB-OD comparisons**.

**S6 Table. ROC analysis results using RT-qPCR with the validation dataset (Related to Fig 3).** 95% confidence intervals are shown, except for AUCs of 1.0. The table is sorted from highest to lowest AUC.

**S7 Table. Log_2_FC estimates, confidence intervals, and Dunnet *P*-values from distinct mycobacterial stimuli in human macrophages *in vitro*.**

**S8 Table. Genes differentially expressed from multibacillary paucibacillary leprosy with |log_2_FC| ≥ 1 and FDR ≤ 0.01.**

**S9 Table. Over-representation analysis (ORA) for MB *vs.* PB (up-regulated) differentially expressed genes.**

**S10 Table. Over-representation analysis (ORA) for MB *vs.* PB (down-regulated) differentially expressed genes.**

**S11 Table. Oligonucleotide sequences.**

## Notes

### Competing Interest Statement

The authors have declared no competing interest.

### Summary of Updates

First round of peer-review. Two authors had their e-mail addresses updated. Figures 1, 2, 3, 4 and 6 have been updated.

