## Appendix 1 for "A new paradigm for leprosy diagnosis based on host gene expression Insights from leprosy lesions transcriptomics" (from another version of this preprint: version 1, 2021-07-30)

### S1 Appendix

### Linking expression profiles to mycobacteria species

*M. tuberculosis* and other mycobacteria-caused diseases are not often differential diagnoses for leprosy, but we sought to investigate how the expression of these genes was modulated by other mycobacterial species. By using publicly bulk RNA-seq and microarray data we were able to compare the expression levels of our gene candidates in two strains of live and inactivated *Mycobacterium tuberculosis*, *M. smegmatis,* and *M. bovis*. Figure 1A shows the expression patterns of 12 available genes (S7 Table). Interestingly, *IDO1* was the most up-regulated gene in *M. smegmatis,* whereas *CXCL10* was the second most induced gene in the same species. Inactivated *M. tuberculosis* H37Rv induced *IDO1* to a lesser extent compared to the other live infections. Some genes were not DE in any of the mycobacteria challenges evaluated, for example *BLK*, *CXCL9*, *MS4A1,* and *TLR10* indicating possible specificity to *M. leprae* response among mycobacteria or cell type specific gene expression. Conversely, *P2RX5* was negatively modulated by *M. smegmatis* only (Fig 1A, S7 Table). Taken together, these results indicate that some specific genes could be used as a specific signature of this mycobacterial infection, and some pathways are probably only induced by *M. leprae* and or they are expressed by other cell types other than macrophages.

In the same manner, we next investigated how *M. leprae* modulated these genes *in vitro*. To this end, the only resource available with live *M. leprae* used Schwann cells, another cell type preferably infected *in vivo*. Schwann cells infected with live *M. leprae* showed some interesting expression patterns for the candidate genes (Fig 1B). *IDO1, APOL3, CXCL10,* and *GBP3* all showed a similar profile of induction only 48 hours post-infection (hpi), where *BIRC3* and *SLAMF7* did the exact opposite. On the other hand, *CD38* showed up-regulation both at 24 and 48 hpi. Finally, *CXCL11* and *CXCL9* both exhibited a similar pattern of repression at 24 and 48 hpi. This result shows that in fact *M. leprae* is capable of modulating these genes *in vitro;* thus, reinforcing the causality between the gene expression phenotype observed in skin biopsies and this pathogen. Also, since *M. leprae* is known to infect Schwann cells in peripheral nerves, these gene expression profiles could also be used to further investigate the disease at that compartment.

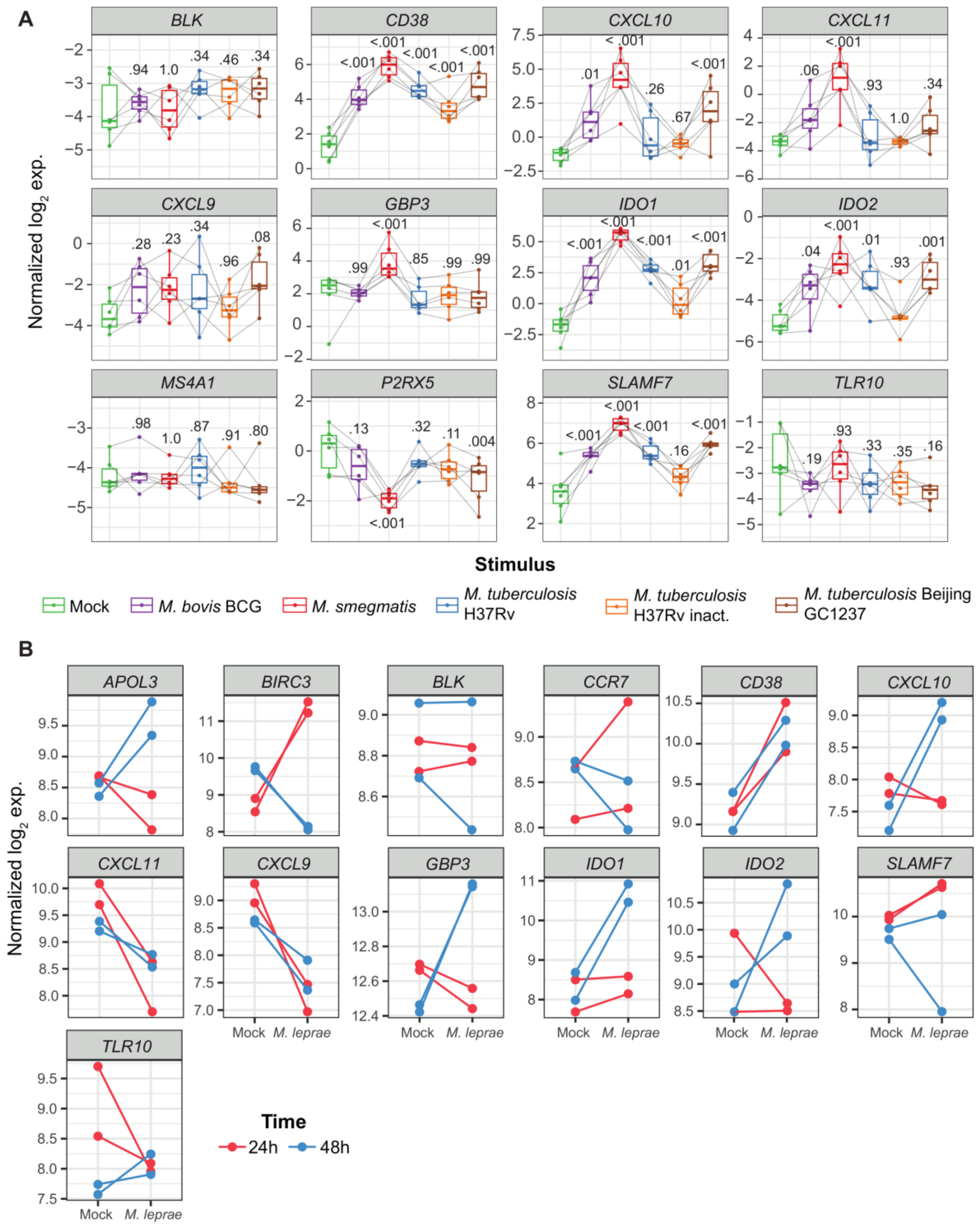


**Fig 1. Gene expression profiles from human macrophages stimulated with mycobacteria *in vitro*.** (A) Box plots showing normalized expression data from monocyte-derived macrophages stimulated with distinct mycobacteria (1:2, MΦ/bacteria) species/strains for 48 hours (GSE67427) [32]. Differences in means from stimulus were compared using gene-wise linear mixed effects with random intercept for each human donor followed by the Dunnet test against the “Mock” group. Numbers on plots are *P*-values. (B) Normalized $\log_2$ expression values from Schwann cells (SC) stimulated with live *M. leprae* (1:10, SC/bacteria) and measured using microarrays (GSE35423) [64]. Related to figure 2.
